# Characterization of new RNA polymerase III and RNA polymerase II transcriptional promoters in the Bovine Leukemia Virus genome

**DOI:** 10.1101/043034

**Authors:** Benoit Van Driessche, Anthony Rodari, Nadège Delacourt, Sylvain Fauquenoy, Caroline Vanhulle, Arsène Burny, Olivier Rohr, Carine Van Lint

## Abstract

Bovine leukemia virus latency is a viral strategy used to escape from the host immune system and contribute to tumor development. However, a highly expressed BLV micro-RNA cluster has been reported, suggesting that the BLV silencing is not complete. Here, we demonstrate the *in vivo* recruitment of RNA polymerase III to the BLV miRNA cluster both in BLV-latently infected cell lines and in ovine BLV-infected primary cells, through a canonical type 2 RNAPIII promoter. Moreover, by RPC6-knockdown, we showed, for the first time, a direct functional link between RNAPIII transcription and BLV miRNAs expression. Furthermore, both the tumor‐ and the quiescent-related isoforms of RPC7 subunits were recruited to the miRNA cluster. We showed that the BLV miRNA cluster was enriched in positive epigenetic marks. Interestingly, we demonstrated the *in vivo* recruitment of RNAPII at the 3’LTR/host genomic junction, associated with positive epigenetic marks. Functionally, we showed that the BLV LTR exhibited a strong antisense promoter activity and provided evidence for a collision between RNAPIII and RNAPII convergent transcriptions. Our results provide new insights into alternative ways used by BLV to counteract silencing of the viral 5’LTR promoter.

## INTRODUCTION

Bovine leukemia virus (BLV) is a B-lymphotropic oncogenic deltaretrovirus infecting cattle that shares common biological and structural features with the human T-cell leukemia virus I and II (HTLV-I and II) (reviewed in (1,2)). In the majority of cases, infection is asymptomatic but 30% of BLV-infected animals will develop a persistent lymphocytosis and less than 5% will progress to B-cell lymphoma or leukemia, termed enzootic bovine leucosis, after a long period of latency characterized by the absence of viral replication (3-5). It is widely accepted that BLV latency is a viral strategy used to escape from the host immune response contributing to tumor development (6). Remarkably, BLV can be experimentally inoculated into sheep that always develop leukemia or lymphoma after a shorter period of incubation than in cattle, and therefore sheep represent a model to study tumor development (7,8).

Transcription of BLV genes initiates at the U3/R junction in the 5’-long terminal repeat (LTR) and is regulated by cellular transcription factors for which several binding sites have been identified in the LTR (9-19), by the viral transactivator TAX_blv_ (20) and by the chromatin status of the BLV provirus (21-25). Indeed, we have previously demonstrated that the 5’LTR RNA polymerase II-driven transcriptional repression is due to the epigenetic state of the 5’LTR characterized by weak histone acetylation and DNA CpG hypermethylation associated to closed chromatin in a lymphoma-derived BLV-infected L267 ovine cell line harboring a fully competent provirus (14,19,21,25).

Recent publications from two independent laboratories have identified, by bioinformatics analysis and deep sequencing, a cluster of 10 micro-RNAs (miRNAs) encoded by the BLV genome (26,27). miRNAs are small ~22 nucleotides RNAs implicated into the regulation of a constantly increasing number of cellular processes and expressed by a wide majority of eukaryotes and some DNA viruses (reviewed in (28)). Despite the 5’LTR silencing dogma, these micro-RNAs are highly transcribed through a noncanonical process, suggesting that the BLV found an alternative way to express a part of its genome in a latency state likely by using an alternative polymerase.

In the present report, we demonstrated the recruitment of a *bona fide* RNA polymerase III at the BLV miRNAs cluster. We also established, for the first time, a direct functional link between BLV miRNAs expression and RNAPIII transcription. In addition, we determined that the promoter responsible for viral miRNAs transcription is a canonical type 2 RNAPIII promoter similar to the one responsible for the tRNAs transcription. Next, we showed that both tumor‐ and quiescent-related isoforms of RNAPIII were recruited to the miRNAs cluster, suggesting that viral miRNAs are produced throughout the complete viral replication cycle, independently of the state of cellular activation. Finally, we demonstrated that the miRNAs cluster exhibited a profile of positive epigenetic marks (histone acetylation and DNA hypomethylation) which is correlated with the high expression of the miRNAs previously reported in BLV-infected cells (27). Interestingly, in addition to the RNAPIII recruitment, we demonstrated an RNAPII recruitment at the junction between the 3’LTR and the host genome. Moreover, this 3’LTR/host region was associated with positive epigenetic marks typical of transcriptionally active promoters. These results prompted us to test the potential promoter activity of the LTR, and we demonstrated that this region was able to drive transcription in the antisense orientation in the nucleosomal context of episomally replicating constructs. This newly discovered RNAPII promoter could be homolog to the HTLV-I 3’LTR antisense promoter responsible for *hbz* transcription (29). Importantly, ChIP-seq experiments performed in L267 cells confirmed the high RNAPIII recruitment to the BLV miRNA cluster and the RNAPII occupancy just downstream of this region suggesting a collision phenomenon between these two polymerase machineries, and stalling of RNAPII.

## MATERIALS AND METHODS

### Cell lines and cell culture

L267 and YR2 BLV infected cell lines were kindly given by Anne Van den Broeke (GIGA-R, University of Liège, Belgium). L267 is a clonal lymphoma-derived B-cell line established from a BLV-infected sheep (S267) injected with naked proviral DNA of an infectious BLV variant (30), whose provirus displays a wild-type sequence (23,31). The L267LT_ax_SN cell line results from the transduction of native L267 with the pLTaxSN retroviral vector expressing the *tax* cDNA following cocultivation with the PG13LT_ax_SN producer cell line and G418 selection of transduced cells (31,32). YR2 is a cloned B-lymphoid cell line established from peripheral blood lymphocytes (PBLs) isolated from a BLV-infected sheep (33), containing a single monoclonally integrated mutated silent provirus (5,32). These cell lines were maintained in Opti-MEM medium (Life Technologies) supplemented with 10% FBS, 1mM sodium pyruvate, 2 mM glutamine, non-essential amino acids and 100μg/ml kanamycin. The 293T cell line (CRL-3216), obtained from American Type Culture Collection (ATCC, Manassas, VA), is established from human embryonic kidney cells that express the large T antigen of simian virus 40 [SV40] (34)). This cell line was maintained in DMEM medium (Life Technologies) supplemented with 10% FBS, 1mM sodium pyruvate and 5% of penicillin-streptomycin. All the cells were grown at 37°C in a humidified 95% air/5% CO_2_ atmosphere. Frozen primary ovine PBMCs were kindly provided by Anne Van den Broeke (Unit of Animal Genomics, GIGA-R, University of Liège, Belgium).

### Plasmid constructs

To construct the episomally replicating contructs, a fragment containing the luciferase gene and the simian virus 40 (SV40) late poly(A) signal was prepared from pGL3-BASIC (Promega) by digestion with KpnI, blunt-ended with DNA polymerase I large (Klenow) fragment, and digestion with BamHI, successively. This fragment was cloned into pREP10s (19) digested with PvuII and BamHI. The resulting plasmid was designed pREP-luc. A fragment containing the complete LTR of the 344 BLV provirus was isolated from the non episomal pLTRwt-luc (10) by digestion with SmaI. This fragment was cloned in both orientations in the pREP-luc digested with BglII and blunt-ended to obtain the pREP-LTR-S-luc and pREP-LTR-AS-luc.

To construct the plasmid expressing the BLV miRNAs, genomic DNA from the L267 cell line was firstly extracted from L267 cells using the DNeasy Blood and Tissue kit (Quiagen) according to the protocol of the manufacturer.

The miRNA cluster was then amplified by PCR using specific primers (Table S2). This PCR reaction was performed at 95 °C for 5 min followed by 35 cycles at 95 °C for 30 sec, 50 °C for 1min 30 sec and by 60 °C for 7min. The obtained product was purified on agarose gel, cloned in the pBS-SKP vector (Promega) and sequenced in order to verify the absence of mutation.

### Chromatin Immunoprecipitation assays

ChIP assays were performed as previously described (25) using the ChIP assay kit (Millipore). Briefly, cells were cross-linked for 10 min at room temperature with 1% formaldehyde and their chromatin were sonicated (Bioruptor Plus, Diagenode) to obtain DNA fragments of 200-400 bp. Chromatin immunoprecipitations were performed with 5 μg of antibodies (see Table S1). Quantitative real-time PCR reactions were performed using the PerfeCTa SYBRgreen SuperMix (Quanta BioSciences). Relative quantification using standard curve method was performed for each primer pair and 96-well Optical Reaction plates were read in a StepOnePlus PCR instrument (Applied Biosystem). Fold enrichments were calculated as percentages of input values or as fold inductions relative to the values measured with IgG. Primer sequences used for quantification (see Table S2) were designed using the software Primer 3.

### ChIP-sequencing assays

ChIP assays were performed as described above. Recovered DNA was then used for library preparation using the TruSeq ChIP Sample Preparation protocol following Manufacter’s instructions (Illumina Technologies). Paired-end sequencing was then performed with the Illumina HiSeq 2000 instrument. More than 20 millions of single reads were obtained for RPC1 and RPB1 libraries and 10 millions of single reads for IgG library. The single-matched reads were mapped to an hybrid ovine genome (OAR_v3.1) containing the BLV provirus sequence (GenBank: KT122858.1) at the L267 integration site (chr10:86299813) using bowtie2 software (-N 1 and ‐k 5 parameters). Finally, our results were visualized using IGV program. Reads spanning LTR-host and LTR-BLV boundaries were extracted from the alignment file using an inhouse R script and ggplot2 was used to plot the ChIP-seq coverages.

### Bisulfite-mediated methylcytosine mapping

Genomic DNA was isolated from L267 and YR2 cell lines using the DNeasy Blood and Tissue kit (Quiagen) according to the protocol of the manufacturer. Genomic DNA was treated with bisulfite with EpiTect Bisulfite kit (Quiagen). The miRNAs cluster region of BLV provirus was amplified by two successive PCR reactions with primers (see Table S2). The first PCR reaction was performed at 95 °C for 5 min followed by 35 cycles at 95 °C for 30 sec, 50 °C for 1min 30 sec and by 60 °C for 7min. The second PCR reaction was performed at 95 °C for 5 min followed by 40 cycles at 95 °C for 30 sec, 50 °C for 1min, 72 °C for 1min 30 sec and by 7 min at 72 °C. The obtained product was purified on agarose gel and cloned in the p-GEMT-easy vector (Promega). At least, 10 clones from each condition were sequenced and analyzed to identify methylated CpGs dinucleotide in this viral region by MethTools software. The frequency of conversion of C to T following sodium bisulfite treatment was greater than 98% at non-CpG sites, indicating the adequacy of the approach.

### Transient transfection and luciferase assays

293T cells were transfected using JetPEI reagent according to the manufacturer’s protocol. 48h post-transfection, cells were collected, lysed and luciferase activities were measured. Firefly luciferase values were normalized with respect to the Renilla luciferase values using the DualGlo-luciferase reporter assay system (Promega).

## RESULTS

### The RNA polymerase III is recruited to the miRNA cluster *in vivo*

Two previous studies have provided indirect evidences that the BLV miRNAs are transcribed by RNA polymerase III (26,27). Indeed RNAPIII initiation cis-elements have been found in the BLV miRNA cluster by bioinformatics analyses (26) and the use of α-amanitin at a concentration known to specifically inhibit RNAPII but not RNAPIII did not decrease BLV miRNA production (26,27). However today, the presence of a functional RNAPIII at the BLV miRNA cluster has not been demonstrated yet. Therefore, we decided to investigate the *in vivo* recruitment to the BLV provirus of RNAPIII by performing chromatin immunoprecipitation (ChIP) assays. For this purpose, we prepared chromatin from the latently BLV-infected L267 cell line and immunoprecipitated the largest RNAPIII subunit (RPC1) using a specific antibody. As controls, we performed ChIP assays with a purified IgG to measure the aspecific background and with an antibody directed against the largest RNAPII subunit (RPB1). Next, purified DNA was amplified by real-time quantitative PCR with oligonucleotide primers hybridizing to twelve specific regions along the BLV genome and its host environment (Fig. 1A). As shown in Figure 1B, we observed recruitment of RPC1 to the BLV miRNA cluster, thereby providing the first *in vivo* evidence that the BLV miRNAs are transcribed by RNAPIII. As positive controls, we amplified the immunopurified DNA by real-time quantitative PCR using oligonucleotide primers hybridizing in genes encoding the transfer RNA (tRNA) for the asparagine amino acid and the 5S ribosomal DNA, both known to be RNAPIII-transcribed genes (reviewed in (35)). Our results showed that RPC1 was recruited to the tRNA^Asp^ and the 5S rDNA genes, demonstrating the functionality of the α-RPC1 antibody we used (Fig. 1E and 1G).

**Figure 1.**
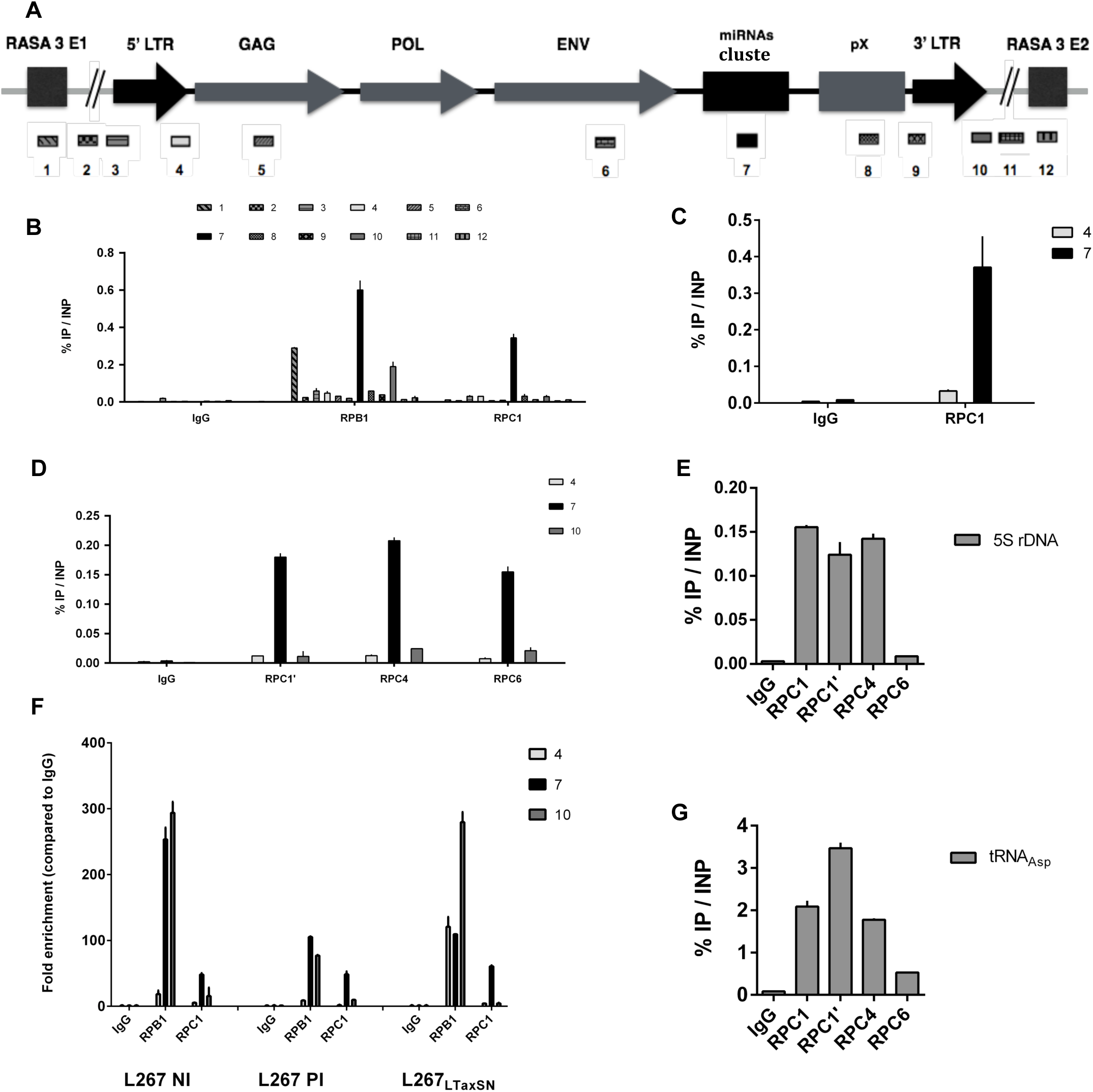
*In vivo* binding of RNAPIII to the BLV miRNA cluster. **(A)** Schematic representation of the BLV provirus in latently-infected L267 ovine cells. The localization of the different amplified regions is presented with respect to the BLV genome. **(B-G)**. Chromatin prepared from L267 cells **(B, D-G)**, BLV-infected ovine PBMCs **(C)** or L267_LTaxSN_ cells **(F)** was immunoprecipitated with specific antibodies directed against different subunits of the RNAPIII as indicated, the largest subunit of the RNAPII or with an IgG as background measurement. Purified DNA was then amplified with oligonucleotide primers hybridizing to either the BLV genome, its host cellular surrounding DNA environment **(B-D and F)** or to the two class III genes encoding 5S rDNA **(E)** and tRNA^asp^ **(G)**. Results are presented as histograms indicating percentages of immunoprecipitated DNA compared to the input DNA (% IP/INP; **B-E and G**) or indicating fold enrichment above the value obtained with IgG, which was arbitrarily assigned the value of 1 **(F)**. Data are the means ± SEM from one representative of at least three independent experiments.

Regarding the recruitment of RNAPII, our previous studies using the L267 cell line failed to demonstrate the recruitment of RNAPII to the 5’LTR, that contains the main transcriptional promoter of BLV genes (21). In the present study, we confirmed these observations and, as a control, we showed the presence of RNAPII to the first exon of the RASA3 gene, the host gene in which the BLV provirus is integrated in the L267 cell line (Fig. 1B). Surprisingly, we observed an important recruitment of RNAPII to the miRNA cluster, in agreement with previously reported ChIP-seq data showing binding of RNAPII near many known RNAPIII genes (36). Interestingly, RNAPII was also recruited at the junction between the 3’LTR and the host genome, suggesting that the 3’LTR may also contains an RNAPII-dependent transcriptional promoter as previously described for the related oncogenic deltaretrovirus HTLV-I (29).

In order to further validate our results, additional ChIP experiments were performed in the YR2 cell line, another latently BLV-infected ovine cell line. These cells are considered as a model for defective latency since two E‐ to K‐ amino acid substitutions in the viral transactivator TAX_BLV_ impair the infectious potential of the integrated provirus (32). Using the YR2 cell line, we observed recruitments of RNAPIII and RNAPII to the BLV genome similar to those observed in the L267 cell line (Fig. S1). However, as opposed to what we observed in the L267 cell line, RNAPII was also recruited at the 5’LTR, confirming our previous results obtained in the YR2 cell line (21) and supporting the deficiency of transactivation mediated by TAX_BLV_ allowing the formation of the RNAPII initiation complex but not an efficient transcriptional elongation (24,37).

To confirm our results in a more physiological model of BLV-infected cells, we next evaluated the *in vivo* recruitment of RNAPIII to the BLV provirus in peripheral blood mononuclear cells (PBMCs) isolated from a BLV-infected sheep that developed leukemia. We demonstrated recruitment of RPC1 to the BLV miRNA cluster (Fig. 1C), thereby confirming, in BLV-infected ovine primary cells, the data we obtained in cell lines. These latter results validated this L267 and YR2 cell lines as an appropriate model to further characterize the RNAPIII promoter present at the BLV miRNA cluster.

To further evaluate the *in vivo* RNAPIII recruitment to the BLV miRNA cluster, we assessed the presence of other RNAPIII subunits by ChIP assays. To this end, sonicated chromatin from L267 cells was immunoprecipitated with specific antibodies directed against different RNAPIII subunits or with a non-specific IgG to measure the aspecific background. Purified DNA was amplified by realtime quantitative PCR with oligonucleotide primers hybridizing to the 5’LTR, the miRNA cluster or the 3’LTR/host junction regions (Fig. 1A). As shown in Figure 1D, we demonstrated that the RPC1, RPC4 and RPC6 RNAPIII subunits were recruited to the miRNA cluster. Therefore, we formally identified the presence of a *bona fide* RNAPIII transcriptional machinery in the BLV miRNA cluster. As positive controls, we showed recruitment of the different RNAPIII subunits to the tRNA^Asp^ and 5S rDNA genes (Fig. 1E and 1G).

Finally, as the BLV 5’LTR is known to be regulated by cellular transcription factors, by the viral transactivator TAX and by the cellular activation state, we tested whether the BLV 5’LTR transcriptional activity could modulate the recruitment of RNAPIII and RNAPII to the BLV miRNA cluster and to 3’LTR/host junction. To this end, we carried out ChIP experiments using chromatin from L267_LTaxSN_ cells, a virus-expressing cell line resulting from the transduction of the L267 cell line with a retroviral vector allowing the overexpression of the viral transactivator Tax_BLV_ (23), or from L267 cells treated with a combination of phorbol 12-myristate 13-acetate (PMA) plus ionomycin, one of the most potent activators of BLV genes expression. Chromatin was sonicated and immunoprecipitated using specific antibodies directed against the largest subunit of RNAPIII and RNAPII or a purified IgG to measure the aspecific background. Purified DNA was then amplified by real-time quantitative PCR with oligonucleotide primers hybridizing to the 5’LTR, the miRNA cluster or the 3’LTR/host junction regions (Fig. 1A). Since the background levels between the two cell lines were different, recruitment of the different proteins was expressed as fold induction compared to the IgG control. Our results demonstrated that, following induction of the 5’LTR transcriptional activity by the PMA/ionomycin cellular treatment (Fig. 1F), recruitment of RPC1 to the miRNA cluster was not affected. In addition, following transactivation of the 5’LTR by overexpression of TAX_BLV_ in the L267_LTaxSN_ cells, we observed no change in the recruitment of the RPC1 subunit to the miRNA cluster nor of the RPB1 subunit to the 3’LTR/host junction (Fig. 1F) compared to the data we obtained in absence of TAX_BLV_ in the L267 cells. As expected, we showed an increase in RNAPII recruitment to the 5’LTR due to the transactivation by TAX_BLV_. On the contrary, we observed a large reduction of RNAPII recruitment to the miRNA cluster in both conditions of activation (see Fig. 1F).

Taken together, our results demonstrate that RNAPIII is recruited to the BLV miRNA cluster *in vivo* and is not dependent on the viral 5’LTR transcriptional state. Moreover, the recruitment of RNAPII occurs at the 3’LTR/host junction, suggesting the presence of a so far unidentified RNAPII promoter in this region.

### BLV miRNAs expression is impaired by depletion of RPC6

Since RNAPIII is recruited *in vivo* to the miRNA cluster, we next investigated a potential functional link between RNAPIII transcriptional machinery and BLV miRNAs expression. To this end, we depleted the RNAPIII subunit RPC6 by RNA interference assays using small hairpin RNAs (shRNAs). With RPC3 and RPC7, the RPC6 subunit has been demonstrated as a member of a subcomplex which is essential for RNAPIII transcriptional initiation but not for RNAPIII transcriptional elongation and termination (38). In addition, according to the literature, RPC6 is the only RNAPIII subunit for which RNA interference experiments have been performed in mammalian cells (39).

To avoid off-target effects of the shRNAs, we used five different shRNAs recognizing distinct target sites in the RPC6 mRNA. We also tested the effect of the combined 5 shRNAs and of a non-targeting shRNA. We first attempted to deplete RPC6 by transfecting or transducing latently BLV-infected L267 and YR2 cell lines. Unfortunately, due to the low transfection or transduction efficiencies in these cell lines, we were unable to show a decrease either in RPC6 protein level or in BLV miRNA transcripts level. Therefore, we performed transient cotransfection experiments in the 293T cell line, using a plasmid expressing the entire miRNA cluster in combination with different shRNAs against RPC6. As an internal control for transfection efficiency, cells were cotransfected with a plasmid expressing the *Firefly* luciferase under the control of the Herpes Simplex Virus thymidine kinase promoter (pTK-luc). Forty-eight hours post-transfection, cells were collected and lysed. Total RNA and protein extracts were used in RT-qPCR and in western blot experiments to measure BLV miRNAs expression levels and RPC6 protein levels, respectively.

Our results showed that all shRNAs directed against the RNAPIII-specific RPC6 subunit decreased expression of BLV-miR-B4-3p (Fig. 2A) and BLV-miR-B2-5p (Fig. 2B), the two most abundant viral miRNAs (27), thereby demonstrating, for the first time, a direct functional link between RNAPIII transcription and BLV miRNAs expression. As controls, we showed that the non-targeting shRNA did not affect BLV miRNAs expression (Fig. 2A and 2B), that the cellular miR-191 transcription, an RNAPII-transcribed miRNA (Li et al, 2014), was not decreased following expression of shRNAs against RPC6 (Fig. 2C) and that transfection efficiencies were similar between the different conditions as shown by the stable *Firefly luciferase* mRNA level (Fig. 2D). Surprisingly, we did not observe any change in RPC6 protein level by western blot, suggesting a low transfection efficiency that did not allow observing any change in RPC6 protein level as only a small fraction of the 293T cellular population was expressing the shRNAs against RPC6.

**Figure 2.**
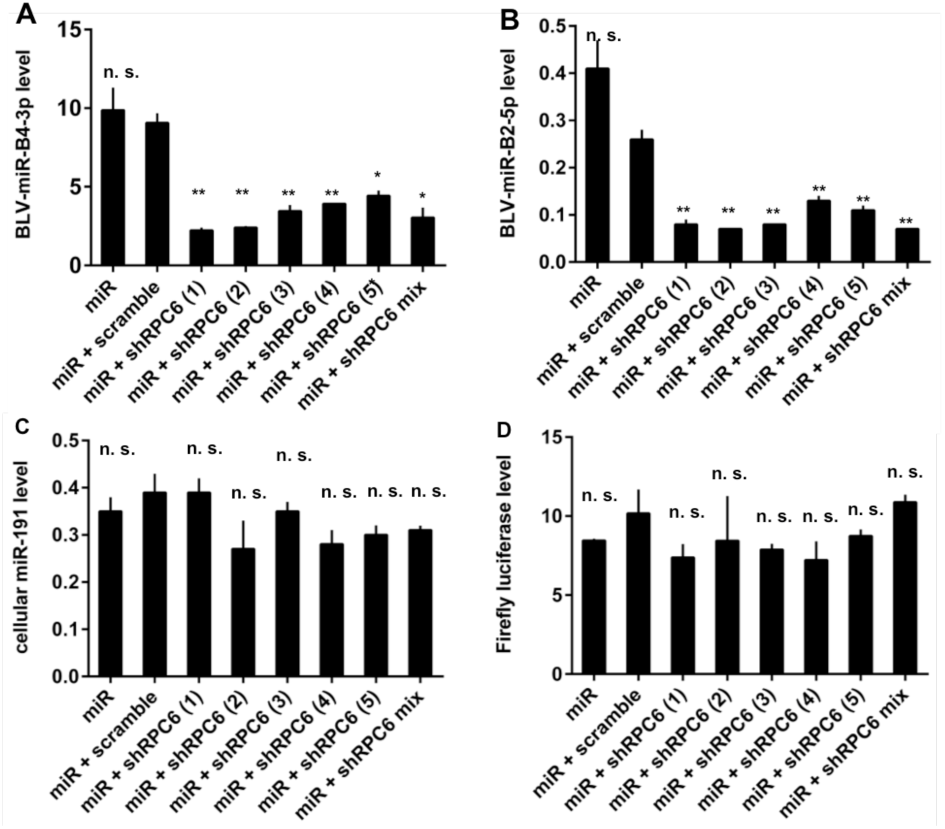
Knockdown of RPC6 induces a decrease of BLV miRNAs expression. The 293T cell line was transiently transfected with a plasmid expressing the miRNA cluster (miR) and plasmids expressing different shRNAs against RPC6 (alone or in a mix) or a non-targeting shRNA. Forty-eight hours posttransfection, total RNA was extracted and assayed for quantitative real-time RT-PCR measuring BLV-miR-B4-3p **(A)**, BLV-miR-B2-5p **(B)**, cellular miR-191 **(C)** or *Firefly* luciferase expression **(D)**. Values were normalized using GADPH gene primers. Data are the means ± SEM from one representative of at two independent experiments.

Thus, our results show that BLV miRNAs expression is impaired following knock-downed expression of RPC6, a member of a subcomplex essential for RNAPIII transcriptional initiation.

### The BLV miRNAs are transcribed from a type 2 RNAPIII promoter

Three different types of RNAPIII promoters are known according to the nature and position of specific *cis*-regulating elements that allow recruitment of specific RNAPIII transcription initiation factors (reviewed in (35,40,41)). Interestingly, each type of promoter is responsible for the transcription of different class III genes: type 1, 2 and 3 promoters are typical of 5S rDNA, tRNAs and small nuclear and nucleolar (i.e U6 RNA and 7SK RNA) genes, respectively.

To further physically characterize the BLV RNAPIII promoter responsible for BLV miRNAs transcription, we assessed the promoter type using ChIP experiments. For this purpose, sonicated chromatin from L267 cells was immunoprecipitated with specific antibodies directed against various RNAPIII transcription initiation factors, or with a purified IgG to measure aspecific background. Purified DNA was amplified by real-time quantitative PCR with oligonucleotide primers hybridizing to the 5’LTR, the miRNA cluster or the 3’LTR/host junction regions (Fig. 1A). As shown in Figure 3A, transcription initiation factors specific for RNA polymerase III were present in the miRNA region of the BLV provirus. More specifically, we did not observe recruitment of TFIIIA, a factor known to be associated only with the type 1 RNAPIII promoter (Fig. 3A). Moreover, we demonstrated the recruitment of TFIIIC, a factor present in both type 1 and type 2 RNAPIII promoters (35). In addition, we showed that the TFIIIB complex present at the BLV miRNA cluster was composed of the Bdp1 and Brf1 subunits in association with TBP, while the Brf2 subunit, specific of the type 3 RNAPIII promoters (35), was absent.

**Figure 3.**
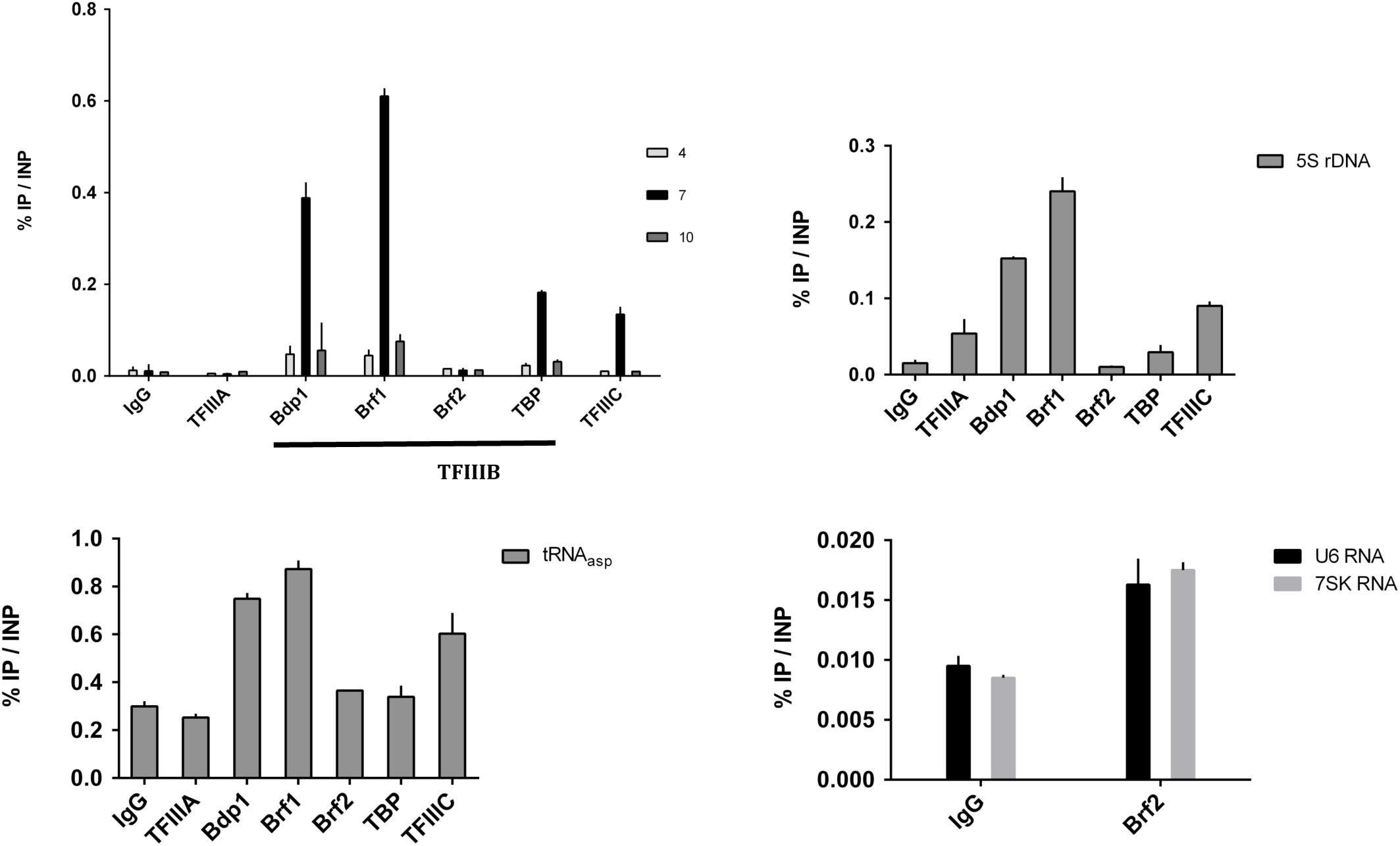
A type 2 RNAPIII promoter drives BLV miRNAs transcription. Chromatin prepared from L267 cells was immunoprecipitated with antibodies directed against different RNAPIII transcription initiation factors or with an IgG as control. Purified DNA was then amplified with oligonucleotide primers hybridizing either to the 5’LTR, the miRNA cluster or the 3’LTR/host junction regions **(A)**, either to the type 1 RNAPIII promoter of the 5S rDNA gene **(B)**, the type 2 RNAPIII promoter of the tRNA^asp^ gene **(C)** or to the type 3 RNAPIII promoter of the U6 and 7SK genes **(D)**. Results are presented as histograms indicating percentages of immunoprecipitated DNA compared to the input DNA (% IP/INP). Data are the means ± SEM from one representative of at least three independent experiments.

In order to validate the specificity of the different antibodies, we amplified immunopurified DNA by real-time quantitative PCR with oligonucleotide primers hybridizing to the 5S rDNA gene containing a type 1 RNAPIII promoter (Fig. 3B), to the tRNA^Asp^ gene containing a type 2 RNAPIII promoter (Fig. 3C) and to the U6 and 7SK genes containing a type 3 RNAPIII promoter (Fig. 3D). Our results showed that all RNAPIII transcription initiation factors were present at the corresponding control regions in accordance with their expected localization, thereby validating the specificity of the antibodies used in ChIP experiments.

Taken together, our results demonstrate that the promoter responsible for transcription of the BLV miRNAs belongs to the type 2 RNAPIII promoter family, as the ones driving tRNAs transcription.

### The two RNA polymerase III isoforms are recruited to BLV the miRNA cluster

Following a duplication event in an ancestor of vertebrates, 2 genes (*POLR3G* and *POLR3GL*) have been found to encode two isoforms of RPC7 (RPC7α and RPC7β, respectively) defining two different RNAPIII complexes (42). The first complex, containing the RPC7α subunit, is restricted to undifferentiated ES cells and to tumor cells, while the second, containing the RPC7β subunit, is ubiquitously expressed and essential for cell growth (43).

In order to investigate whether cell transformation was required for BLV miRNAs transcription, we evaluated the *in vivo* recruitment to the viral miRNA cluster of the two different RNAPIII complexes. To this end, we performed ChIP assays using chromatin obtained from L267 cells and antibodies directed against the RPC7α and RPC7β subunits or a purified IgG to measure aspecific immunoprecipitation background. Next, we amplified immunopurified DNA by real-time quantitative PCR with oligonucleotide primers hybridizing to the 5’LTR, the miRNAs cluster or the 3’LTR/host junction regions (Fig. 1A). As shown in Figure 4A, we showed that both RPC7a and RPC7b subunits were recruited to the BLV miRNAs cluster.

**Figure 4.**
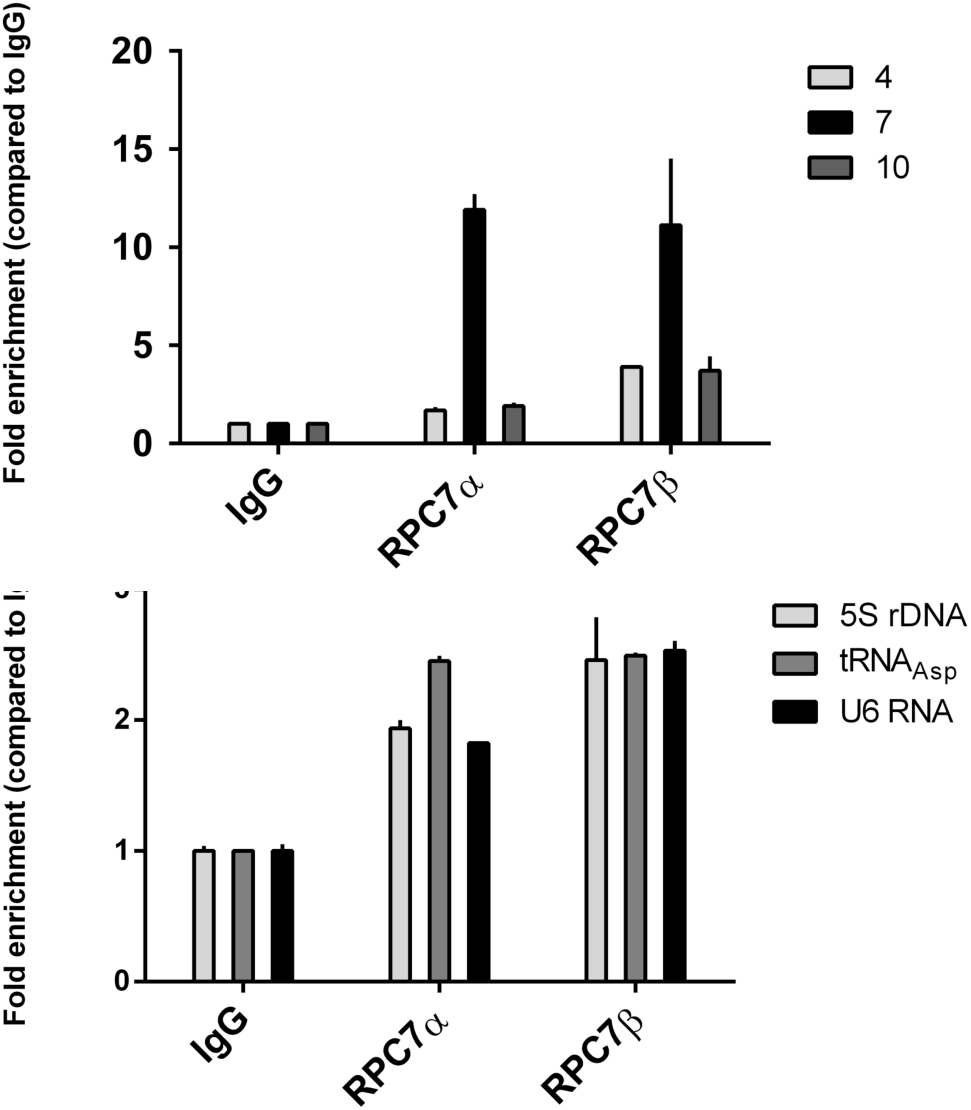
Recruitment of the RPC7α and RPC7β subunits to the BLV miRNA cluster. Chromatin prepared from L267 cells was immunoprecipitated with specific antibodies directed against the two isoforms of the RPC7 subunit or with an IgG as background measurement. Purified DNA was then amplified with oligonucleotide primers hybridizing to the 5’LTR, the miRNA cluster or the 3’LTR/host junction regions **(A)** or to class III control genes **(B)**. Results are presented as histograms indicating fold enrichments compared to the value obtained with IgG to which a value of 1 was assigned. Data are the means ± SEM from one representative of at least three independent experiments.

As positive controls, we amplified immunopurified DNA by real-time quantitative PCR with oligonucleotide primers hybridizing in three known RNAPIII-dependent genes. As shown in Figure 4B, both RPC7α and RPC7β subunits were recruited to these three genes. Similar experiments were performed with sonicated chromatin from the YR2 cell line (Fig. S2). Interestingly, as opposed to what we observed in the L267 cell line, only the RPC7a subunit appeared to be recruited to the BLV miRNA cluster and to control RNAPIII-dependent genes in the YR2 cell line, while both subunits are expressed in YR2 cells (Anne Van den Broeke, personal communication). This unexpected result could possibly be explained by a differential recruitment of RPC7α and RPC7β, after cellular transformation, in the L267 cell line compared to the YR2 cell line.

Taken together, our results suggest that the two different isoforms of RNAPIII are able to transcribe the BLV miRNAs at all stages of the BLV disease, in agreement with previous RNA-seq data showing miRNAs expression in transformed and non-transformed cells (27).

### The BLV miRNA cluster DNA is not methylated

It is known that CpG islands, DNA regions where enriched CpG dinucleotides are observed, may modulate transcriptional activity of a nearby promoter, according with their methylation status (44). In addition, our laboratory has previously demonstrated that DNA methylation is an important epigenetic chromatin modification that represses BLV transcription from the 5’LTR in the true latent L267 cell line (25). In order to determine whether the BLV genome contains CpG islands, we performed *in silico* analyses using three different prediction tools (Methprimer, CpGPlot and CpG Island Searcher) that all identified a CpG island containing the BLV miRNA cluster and spanning a region from nucleotide (nt) 6342 to nt 6833 (nt +1 defined as the first nucleotide of the 5’LTR after retrotranscription). Therefore, we decided to examine the role of DNA methylation as a potential epigenetic regulator of BLV miRNAs expression. To this end, we evaluated the DNA methylation status of each CpG dinucleotide of the BLV miRNAs cluster in the context of integrated proviruses present in L267 and YR2 cells by treating genomic DNA with sodium bisulfite. Our results showed that, except for the third and the fourth CpG located upstream of the first BLV pre-miR, the miRNA cluster were unmethylated in both the L267 (Fig. 5) and YR2 (Fig. S3) cell lines.

**Figure 5.**
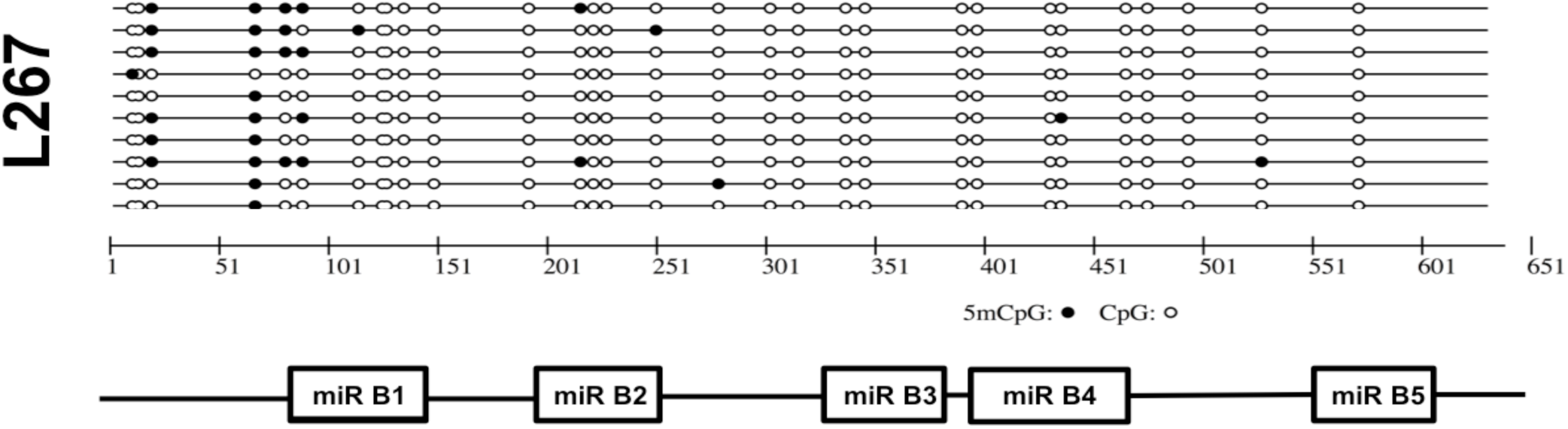
Evaluation of the CpG dinucleotide methylation status of the BLV miRNA cluster. Genomic DNA from L267 cells was extracted and treated with sodium bisulfite and the miRNA cluster was amplified by nested PCR. These amplified products were subcloned in a TA cloning vector and 10 independent clones were sequenced. Open and filled circles represent unmethylated and methylated CpG dinucleotides, respectively. The position of the 5 BLV-pre-miRs with respect to the miRNA cluster is presented in the lower part of the figure.

In conclusion, our results indicate that the BLV miRNA cluster exhibits an hypomethylated CpG profile, in agreement with the high expression level of the BLV miRNAs (27).

### Epigenetic marks associated with active promoters are present to the miRNA cluster and 3’LTR/host cellular DNA junction

In order to further characterize the epigenetic environment present at the BLV miRNA cluster and 3’LTR/host cellular DNA junction, we performed ChIP assays using chromatin from L267 cells and antibodies directed against histone post-translational modifications or the H2A.Z histone variant. As control, we performed ChIP assays with a purified IgG to measure aspecific immunoprecipitation background. Purified DNA was then amplified by real-time quantitative PCR with oligonucleotide primers hybridizing to the 5’LTR, the miRNA cluster or the 3’LTR/host cellular DNA junction regions (Fig. 1A). As shown in Figure 6A, we observed the presence, at the level of the miRNA cluster, of acetylated histone H3 (H3Ac), acetylated histone H3 lysine 9 (H3K9Ac) and trimethylated lysine 4 histone H3 (H3K4me3), three chromatin marks associated with RNAPII‐ and RNAPIII active promoters (45,46). Interestingly, those marks were also present at the 3’LTR/host cellular DNA junction, further supporting the presence of an active promoter in this region. In agreement with these results, we did not detect H3K9me3 and H3K27me3, two repressive epigenetic marks associated with transcriptional silencing. As positive controls, we showed the presence of these two repressive marks at the *ZNF12* and *HOXD10* promoter regions (Fig. 6B) as previously reported (47,48).

**Figure 6.**
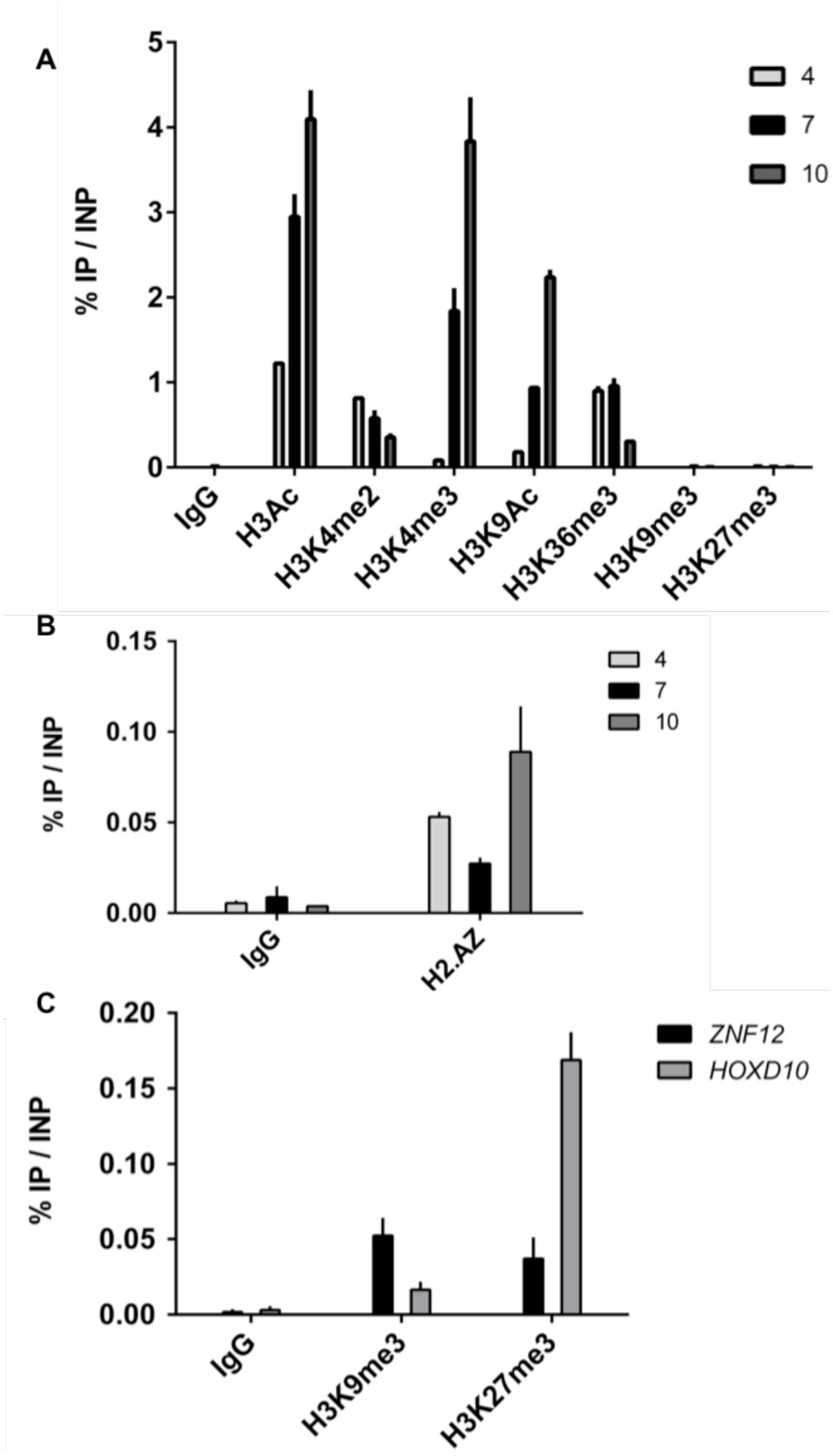
The BLV miRNA cluster and 3’LTR/host genome junction are associated with active epigenetic marks. Chromatin prepared from L267 cells was immunoprecipitated with specific antibodies directed against different histone post-translational modifications **(A,C)**, the H2A.Z histone variant (**B**) or with an IgG as background measurement. Purified DNA was then amplified with oligonucleotide primers hybridizing to the 5’LTR, the miRNA cluster or the 3’LTR/host junction regions (**A** and **C**) or to the promoter regions of the *ZNF12* and *HOXD10* genes (**B**). Results are presented as histograms indicating percentages of immunoprecipitated DNA compared to the input DNA (% IP/INP). Data are the means ± SEM from one representative of at least three independent experiments.

Interestingly, our results showed the presence of H3K4me2 and the histone H2A.Z variant, two epigenetic marks associated with active promoters and enhancers (46) in the three tested regions (Fig. 6A and 6C). Moreover, we observed the presence in the three regions of H3K36me3, a positive epigenetic mark associated with RNAPII transcriptional elongation. Remarkably, since the level of H3K36me3 has been previously reported to increase within gene bodies (46) and since the profile of H3K36me3 shown here was increasing from the 3’LTR to the 5’LTR, our results were in agreement with the presence of an antisense transcription starting from the 3’LTR.

Together, our data strongly support the presence of active promoters at the BLV miRNAs cluster and at the junction between the 3’LTR and the host genomic region.

### The region of RNAPII recruitment in the BLV 3’LTR exhibits antisense promoter activity

The *in vivo* recruitment of RNAPII at the 3’LTR/host junction (Fig. 1B) and its epigenetic profile characteristic of an active promoter (Fig. 6A and 6C) prompted us to test its potential promoter activity in a nucleosomal context. To this end, we decided to use pREP-based episomal reporter constructs containing the EBV replication origin and encoding nuclear antigen EBNA-1. Indeed, several studies have reported that pREP-based episomal constructs display hallmarks of proper chromatin structure when transiently transfected into cells (49,50). Thus, we cloned the complete BLV LTR upstream of the luciferase (luc) gene in both the 5’ and 3’ orientations into a modified pREP10 episomal vector (pREP-luc). The episomally replicating BLV LTR-luc we generated (named pREP-LTR-S-luc and pREP-LTR-AS-luc, respectively) were transfected into the human B-lymphoid Raji cell line. Forty-hours post-transfection, cells were lysed and assayed for luciferase activity (Fig. 7). The pREP-LTR-S-luc and the pREP-LTR-AS-luc constructs presented a 42‐ and a 357-fold increases in luciferase activity compared to the parental reporter vector pREP-luc. Therefore, our results are consistent with the fact that the BLV 3’LTR exhibits an antisense promoter activity and that this transcriptional activity is significantly higher than the 5’LTR sense promoter activity which is responsible for BLV mRNAs transcription.

**Figure 7.**
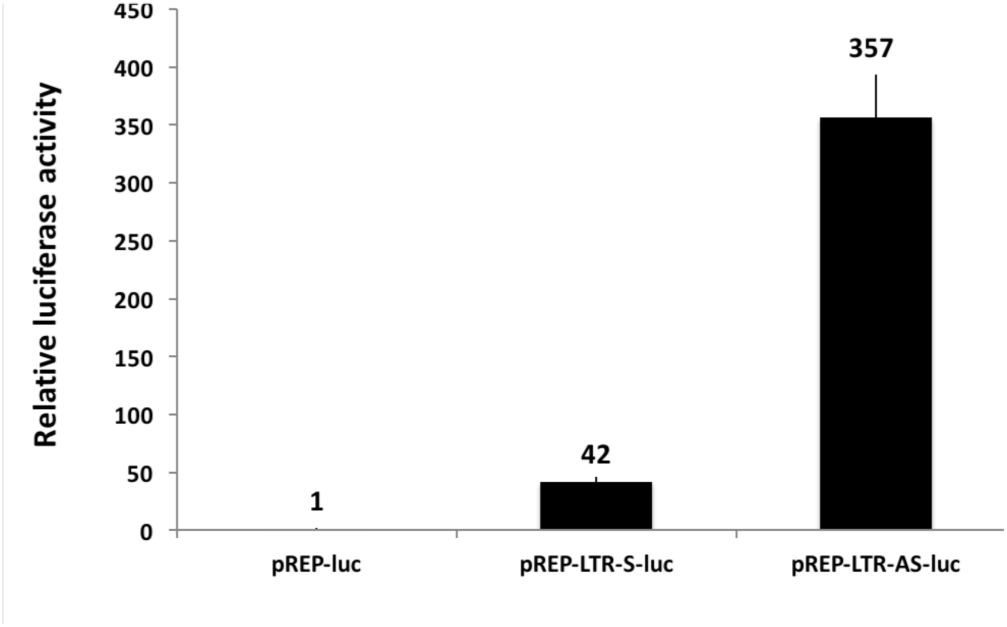
The RNAPII promoter in the 3’LTR is transcriptionally active. Raji cells were transiently transfected with 400ng of pREP-luc, pREP-LTR-S-luc or pREP-LTR-AS-luc constructs and 20ng of pRL-TK. 48h after transfection, cells were collected, lysed and both firefly and renilla luciferase activities were measured. Results are presented as histograms indicating relative luciferase activities compared to the pREP-luc for wich a value of 1 was assigned. Data are the mean ± SEM from one representative of at least three independent experiments.

Taken together, our results demonstrate the presence of a transcriptionally active RNAPII promoter in the BLV 3’LTR, associated with positive epigenetic marks and responsible for antisense transcription, a situation reminiscent of the previously reported HTLV-I 3’LTR antisense promoter activity responsible for transcription of the viral *hbz* gene (29).

### The RNAPIII collides the RNAPII at the BLV miRNA cluster

As presented in Figure 1B, we showed an unexpected recruitment of RNAPII to the BLV miRNA cluster. As this presence could be explained by a cooperative binding (36) or by the antisense transcription coming from the 3’LTR (Fig. 7), we decided to investigate the RNAPII and RNAPIII coverage along the BLV genome by ChIP experiments followed by deep sequencing analysis. To this end, sonicated chromatin from L267 cells was immunoprecipitated with specific antibodies directed against the largest subunits of RNAPII (RPB1) and RNAPIII (RPC1) or with a purified IgG as background measurement. The immunoprecipitated DNA was then used for paired-end deep sequencing analysis. Reads were aligned to the ovine reference genome, where BLV sequence has been inserted at the integration site of the L267 cell line. Results of single-matched are presented in Figures 8A.

**Figure 8.**
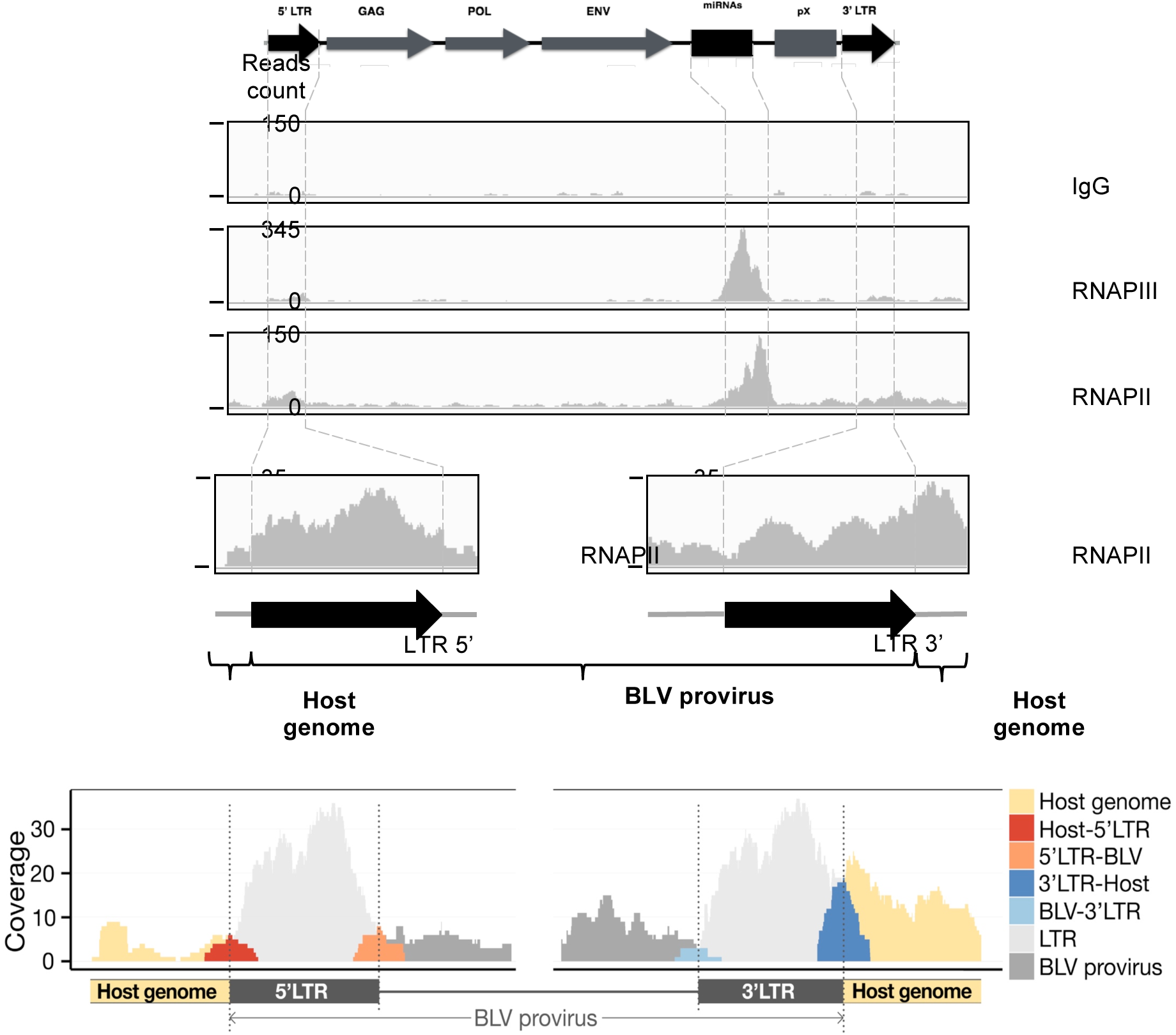
Collision between RNAPIII and RNAPII at the BLV miRNA cluster. View of the tags obtained following ChIP experiments using chromatin prepared from L267 cells and immunoprecipitated with specific antibodies raised against the largest subunit of the RNA polymerase II (RPB1), the largest subunit of the RNA polymerase III (RPC1) or with a purified IgG as background measurement. **(A)** Single matched reads for RNAPII/RNAPIII coverage along the BLV genome. **(B)** Zoom-in of the 5’ and 3’LTRs regions for single and double matched reads.

As shown in Figure 8A, we confirmed the presence of both RNAPII and RNAPIII complexes at the BLV miRNA cluster in agreement with our quantitative PCR results (Fig. 1B). Interestingly, our results revealed the presence of RNAPIII at the miRNA cluster, while RNAPII coverage was further downstream. This localization of RNAPII downstream relative to the RNAPIII transcriptional start site contrasted with previous studies showing cooperation between these two polymerases when RNAPII is present upstream the RNAPIII-dependent unit (45,51). Therefore, our ChIP-seq data supports the evidence of an antisense RNAPII-dependent transcription initiating in the 3’LTR. Moreover, there was a drastic drop in RNAPII ChIP-seq coverage at the miRNA cluster downstream of the RNAPIII coverage, further supporting convergent transcriptions and collision between RNAPIII and RNAPII (Fig. 8A). However, no recruitment of RNAPII to the 3’LTR was observed. This result was due to the low quality of the read alignment in the LTRs and explained by the formation process of these LTRs during the reverse transcription leading to regions with identical nucleotidic sequences therefore avoiding unambiguous mapping of the reads. In order to have a better view of the RNAPII coverage at the different LTRs, we performed a new alignment of the reads allowing one or two matches to our reference genome. We decided to focus on unique DNA sequences comprising the regions adjacent to the LTRs boundaries (Figure 8B). These results revealed a higher coverage of the 3’LTR/host junction, (22 and 2 reads covering the junction after chromatin immunoprecipitation with RNAPII antibody and IgG, respectively) compared to that of the 5’LTR/viral genome junction (8 and 5 reads covering the junction after chromatin immunoprecipitation with RNAPII antibody and IgG, respectively). However, we show that the coverage of the 3’LTR/viral genome (5 and 0 reads covering the junction after chromatin immunoprecipitation with RNAPII antibody and IgG, respectively) and of the 5’LTR/host genome (5 and 1 reads covering the junction after chromatin immunoprecipitation with RNAPII antibody and IgG, respectively) were similar.

Altogether, our data are further confirming RNAPII recruitment to the transcriptional promoter located at the end of the 3’LTR (Fig. 8B) and indicate the presence of an active RNAPII-dependent promoter in the BLV 3’LTR, responsible for an antisense transcription and colliding with the RNAPIII transcribing the viral miRNA cluster.

## DISCUSSION

One of the major features of BLV infection is the absence of viremia due to the RNA polymerase II 5’LTR-driven transcriptional repression. Nevertheless, recent reports using bioinformatics analysis (26) and high throughput sequencing of small RNAs from BLV-infected cells (27) have shown the presence of a cluster coding for viral miRNAs located between the end of *env* gene and the pX region. Some indirect evidence support the notion that the miRNA cluster is transcribed by the RNA polymerase III, although a direct functional link between RNAPIII transcription and BLV miRANs expression has not been reported to date. Indeed, the algorithm used by Kincaid and coworkers has been designed to find RNAPIII-specific initiation *cis*-regulatory elements in the genome of retroviruses and this analysis has identified BLV as the only retrovirus containing such *cis*-regulatory elements (26). Moreover, α-amanitin treatment at a concentration known to inhibit specifically RNAPII but not RNAPIII transcriptional elongation (52) did not affect BLV miRNA expression (26,27). In the present report, we aimed at providing a direct demonstration that the RNAPIII is responsible for BLV miRNA transcription and is recruited to the viral miRNA cluster *in vivo*, and at epigenetically and functionally characterizing its promoter. To this end, we performed chromatin immunoprecipitation assays in two BLV latently-infected ovine cell lines and in BLV-infected primary B cells.

We firstly demonstrated that all tested RNAPIII subunits were recruited to the miRNA cluster *in vivo* (Fig. 1B and 1D), supporting the presence of a *bona fide* RNAPIII at this genomic region and confirming previous evidences (26,27). Furthermore, we also demonstrated the recruitment of the RPC1 RNAPIII subunit in primary PBMCs from a BLV-infected leukemic sheep suffering from leukemia (Fig. 1C), confirming our results in a more physiological model of BLV infection. We also showed that RNAPIII recruitment to the miRNA cluster was not affected by induction of 5’LTR-dependent viral expression either by ectopic expression of TaxBLV (L267LTaxSN cell line) or by PMA/ionomycin treatment (Fig. 1F). These data suggest that the level of 5’LTR transcriptional activity does not seem to influence the recruitment of RNAPIII, in agreement with the stable level of miRNA expression previously observed in L267_LTaxSN_ cells compared to L267 cells (27). After confirming the *in vivo* recruitment of RNAPIII to the BLV miRNA cluster, we showed that the viral miRNA expression was significantly decreased upon depletion of RPC6 (Fig. 2A and 2B), a specific RNAPIII subunit implicated in RNAPIII transcriptional initiation (38), functionally demonstrating for the first time that the BLV miRNAs are transcribed by RNAPIII. Moreover, two forms of mammalian RNAPIII (α or β) have been identified depending on the presence of RPC7α or RPC7b subunit isoforms, respectively (42,43). While the RNAPIIIb is essential for undifferentiated and differentiated cells, the RNAPIIIa is mainly present in undifferentiated and tumor cells, and the level of the RPC7α mRNA increases during cellular transformation (43). Interestingly, we showed here that both isoforms of RPC7 were recruited to the miRNA cluster (Fig. 4), suggesting that BLV miRNAs are transcribed by the two forms and do not require cellular transformation. Our results are in agreement with previously reported RNA-seq data (27).

Compared to the promoter complexity of RNAPII transcribed genes, the promoter driving RNAPIII transcribed genes is much more simple. Indeed, RNAPIII promoters are composed of a relatively small number of cis-acting elements that are generally located within the transcriptional unit, and regulated by a small number of transcription factors (reviewed in (40)). In this study, we showed that the TFIIIA initiation factor, which is specific of type 1 RNAPIII promoter driving transcription of the 5S rDNA, was not recruited to the BLV miRNA cluster. However, we demonstrated that the TFIIIB (composed of TBP, BDP1 and BRF1) and TFIIIC complexes were present at the miRNA cluster, supporting the use of a canonical type 2 RNAPIII promoter for BLV miRNA transcription (Fig. 3A) as for cellular tRNA transcription. These results confirm the prediction of the algorithm, based on cis-regulatory elements, used by Kincaid and coworkers (26).

Our laboratory has previously demonstrated that DNA methylation and histone tail post-translational modifications are important epigenetic chromatin modifications regulating BLV gene expression (19,21,24,25). Indeed, we have previously reported that 5’LTR hypermethylation is associated with the BLV repression in the true latent L267 cell line but not in the defective latency state present in the YR2 cell line (25). Here, we showed by bisulfite genome sequencing that the DNA encoding the BLV miRNAs cluster, which is located within a CpG island, was not methylated. To further characterize the epigenetic landscape associated with the miRNA cluster, we demonstrated the absence of repressive epigenetic marks (H3K9me3 and H3K27me3) and the presence of activating epigenetic marks associated with RNAPII‐ and RNAPIII-dependent active promoters (45,46). More specifically, we showed the presence of acetylated histone H3 (H3Ac, H3K9Ac), di-and tri-methylated lysine 4 histone H3 (H3K4me2, H3K4me3). Overall, the activating chromatin environment of the miRNA cluster we demonstrated in the BLV latently-infected cell lines were consistent with the high level of expression of the miRNAs observed in RNA-seq experiments (27).

The ChIP results presented here showed an *in vivo* recruitment of RNAPII at the 3’LTR/host junction (Fig. 1B). This specific recruitment was associated with the presence of positive epigenetic marks (such as histone acetylation, H3K4me2, H3K4me3 and the H2A.Z histone variant) and the absence of repressive marks (such as H3K9me3, H3K27me3 (Fig. 6A) or DNA methylation (25)). Moreover, using episomally replicating luciferase reporter constructs that display the hallmarks of proper chromatin structure when transiently transfected into cells (49,50), we demonstrated that the BLV 3’LTR exhibited an antisense promoter activity. Such a promoter at the end of the BLV 3’LTR could be the counterpart of the previously reported antisense RNAPII promoter responsible for the transcription of the *hbz* gene in the closely related HTLV-1 deltaretrovirus (29). Of note, HbZ expression is important for the progression of adult T-cell leukemia since it favors proliferation of HTLV-1-infected cells (53). BLV 3’LTR antisense promoter activity reported here is further supported by the presence of H3K36me3 along the BLV provirus that is lower at the 3’LTR/host junction than at the miRNA cluster and the 5’LTR. Indeed, H3K36me3 level generally increases along RNAPII transcribed regions (46), whereas RNAPIII transcribed genes lack for enrichment of this epigenetic mark (45). In addition, RNA-seq experiments of primary tumors and pre-leukemic clones described by our collaborators Durkin and co-workers, reported on the bioRxiv preprint server (54), revealed the production of two antisense transcripts originating in the BLV 3’LTR. Their identification in animals and characterization are described in this latter manuscript.

It has been shown that α-amanitin treatment, an inhibitor of RNAPII transcriptional elongation when used at low concentration (52) does not decrease BLV miRNAs expression (26,27), thereby indicating that RNAPII is not responsible for the BLV miRNAs transcription. Nevertheless, we showed here the recruitment of RNAPII to the miRNA cluster (Fig. 1B). It is worth noting that Barski and coworkers have previously shown the presence of RNAPII at RNAPIII-transcribed *loci* by ChIP-seq experiments (45). In addition, it has been proposed that RNAPII activity might facilitate the expression of some RNAPIII products by modifying the local chromatin structure (51). However, in the context of the BLV provirus, our ChIP-seq experiments suggest that the presence of RNAPII at the miRNA cluster is the result of a collision phenomenon between RNAPIII transcribing the BLV miRNAs in the sense orientation and RNAPII transcribing from the 3’LTR in the antisense orientation (Fig. 8A and 8B). It has been previously demonstrated *in vitro* that collided RNAPII polymerases resulting from convergent transcription can stop without dissociation from the genomic unit (55). A similar phenomenon could therefore explain the *in vivo* RNAPII recruitment we observed here in our ChIP and ChIP-seq experiments (Fig. 1B and 8A). Nevertheless, to date, the biological role of convergent transcriptions remains largely unknown. Some studies have shown that antisense transcription could be involved in the masking of splicing sites and thus responsible for the production of different transcript isoforms (56), whereas other studies have shown that sense and antisense transcriptions can be parts of a self-regulatory circuit allowing a finer-tuned regulation between their respective expression (57). In the context of the BLV provirus, at the miRNA cluster, RNAPII could be responsible for the regulation of miRNAs expression or alternatively, as suggested by the two antisense transcript isoforms described in the manuscript of Durkin and co-workers (54), the RNAPIII could allow an alternative splicing of the BLV antisense transcripts.

In conclusion, our data demonstrated for the first time a direct functional link between RNAPIII transcription and BLV miRNAs expression, and the *in vivo* recruitment of a *bona fide* RNAPIII complex to the BLV miRNA cluster through a type 2 RNAPIII promoter in BLV-latently infected cell lines and ovine BLV-infected primary cells. Moreover, we demonstrated that the BLV LTR was able to drive transcription in the antisense orientation, as previously reported for the closely related HTLV-1 retrovirus. Importantly our ChIP and ChIP-seq experiments demonstrated a high RNAPIII recruitment peak in the BLV micro-RNA cluster and a RNAPII recruitment peak just downstream of this region, suggesting a collision phenomenon between the two polymerases and stalling of RNAPII. The two new RNAPIII and RNAPII transcriptional promoters located in the BLV genome brings additional factors in an already complex network of regulators affecting the level of BLV expression and potentially deltaretrovirus induced leukemia.

## FUNDING

This work was supported by grants from the Belgian *Fonds de la Recherche Scientifique* (FRS-FNRS, Belgium), the Télévie program of the FRS-FNRS and the Van Buuren and Jean Brachet Foundations. CVL is Directeur de Recherches of FRS-FNRS. BVD is a postdoctoral fellow from Walloon region excellence program “CIBLES”. AR is a doctoral fellow from the Belgian *Fonds pour la formation à la Recherche dans l’Industrie et dans l’Agriculture* (FRIA). ND is supported by a « PDR » grant from the FRS-FNRS and SF is a post-doctoral fellow from the Télévie program.

## ACKNOWLEDGMENT

We acknowledge A. Van den Broeke, N. Hernandez, R. White for generously providing reagents used in this work. We acknowledge M. Defrance (Erasme Hospital, ULB, Belgium) and N. Rosewick (GIGA-R, University of Liège, Belgium) for their help in the bioinformatical processing of our ChIP-seq data. We acknowledge A. Van den Broeke for critical reading of the manuscript. The authors wish it to be known that, in their opinion, the last two authors should be regarded as joint senior authors.

